# Insights into the transmission of seedborne *Pseudomonas syringae* strains that cause zucchini diseases

**DOI:** 10.64898/2026.08.21.746309

**Authors:** Caroline Lacault, Marie-Agnès Jacques, Armelle Darrasse

## Abstract

Vein clearing of zucchini (VCZ) and bacterial leaf spot (BLS) are caused by various strains of the *Pseudomonas syringae* species complex that infect zucchini (*Cucurbita pepo*) seeds. VCZ strains have a narrow host range of cucurbits and affect only seedlings, whereas BLS strains have a broader host range and cause symptoms on adult plants. A multiplex qPCR test showed that VCZ strains predominated in infected seed lots produced in different countries. We surveyed hybrid seed crops grown in parallel in two French regions to address inoculum sources. According to DNA-based approach, parental seed lots were positive to BLS strains, although no culturable bacteria were recovered. Hybrid seed lots produced in the Rhone Valley (southeastern France) showed higher infection rates than those produced in Limagne (central France), and VCZ strains were recovered only from the Rhone Valley. Two representative strains of VCZ and BLS colonized seeds through the vascular and floral pathways, whereas only the BLS strain was transmitted through the pericarp. These findings suggest that floral transmission, potentially mediated by pollinators, could explain the predominance of VCZ strains under favorable regional conditions, and that pericarp transmission in BLS strains could be linked to their capacity to cause disease on adult plants. Furthermore, some infections undetected in seeds became apparent after germination, indicating that testing germinated seeds rather than seeds could help seed industry to take in account only bacterial infections transmitted to the seedling. Together, these results provide valuable insights into the epidemiology of *P. syringae* transmission to zucchini seeds.

## Introduction

Since the early 2000s, the zucchini seed industry has been facing a bacterial disease, Vein Clearing of Zucchini (VCZ), which is transmitted through seeds. VCZ is damaging to seedlings, although no outbreaks have been reported on mature plants (Lacault et al. 2020; Manceau et al. 2011). The seedlings suffer from stunted growth, which can lead to a complete growth arrest. Necrotic spots on the cotyledons and deformities, such as twisting and thickening of the stem base, are sometimes observed. When the seedling survives, necrosis and deformities may spread to the first leaves and be accompanied by a temporary symptom of vein clearing, a characteristic that gave this disease its name. This disease is particularly prevalent in cool humid growing conditions.

VCZ symptoms on seedlings are caused by various strains of the *Pseudomonas syringae* species complex, which are clearly distinct from other pathovars pathogenic on cucurbits, such as *P. syringae* pv. *aptata* and *P. syringae* pv. *lachrymans* (Bradburry 1986; Lacault et al. 2020; Manceau et al. 2011). While the strains isolated during the early outbreaks were highly homogeneous, a larger diversity of strains has been isolated more recently from seed lots and seedlings. Indeed, some strains now share high average nucleotide identities based on BLAST (ANIb) values with bacterial leaf spot (BLS) strains that have caused outbreaks in adult cucurbit plants in the United States and Australia. Combined with the results of pathogenicity tests, this observation indicates that strains isolated from zucchini seeds and have a wide host range of cucurbits are related to BLS strains (Djitro et al. 2022a; b; Lacault et al. 2020; Newberry et al. 2016; 2017; 2018; 2019; Willmann et al. 2026). All these strains form several phylogenetic clusters, that are distributed across two clades: clade 2a corresponding to *Pseudomonas cerasi* (Berge et al. 2014, Gomila et al. 2017) and clade 2ba, resulting from recombination events between *P. cerasi* and *P. syringae* (Newberry et al. 2019). Comparative genomics of type three effector (T3E) repertoires distinguishes two main pathogenic groups within these clusters: strains carrying *avrRpt2* having a narrow host range restricted to *Cucurbita* spp., and strains carrying *hopZ5* having a broader host range that also includes *Cucumis melo*, *Cucumis sativus*, and sometimes *Citrullus lanatus* (Djitro et al. 2022b; Lacault et al. 2020; Newberry et al. 2019). The *avrRpt2*^+^/*hopZ5*^-^ genotype corresponds to the historical strains first associated with VCZ outbreaks on seedlings, informally described as the “*peponis*” pathovar, which has never been reported on adult plants (Lacault et al. 2020; Manceau et al. 2011). The *avrRpt2*^-^/*hopZ5*^+^ genotype, by contrast, includes the highly virulent strains responsible for BLS outbreaks on commercial watermelon, cantaloupe, and squash in Florida (Newberry et al. 2019). For simplicity, we hereafter refer to the narrow host range strains (*avrRpt2*^+^/*hopZ5*^-^) as VCZ strains and to the broad host range strains (*avrRpt2*^-^ /*hopZ5*^+^) as BLS strains (Table 1). Strains belonging to two additional clades, 2b (*P. syringae sensu stricto*) and 2d, are also isolated from zucchini seeds, but do not appear associated with outbreaks (Lacault et al. 2020; 2024), consistent with a recent study showing their lower pathogenicity and prevalence on zucchini relative to dominant clades 2ba (78%) and 2a (13%) found in commercial seed production (Willmann et al. 2026).

**Table 1.** qPCR identification profiles for *Pseudomonas syringae* strains isolated in zucchini seed lots.

| MLST group<sup>1</sup> | detection of qPCR marker<sup>2</sup> |  |  |  | identification<sup>3</sup> |
| --- | --- | --- | --- | --- | --- |
|  | *sylC* | *avrRpt2* | *hopZ5* | clade 2d |  |
| 2ba-A | + | + | - | - | VCZ |
| 2ba-C1 | + | + | - | - | VCZ |
| 2ba-B | + | - | + | - | BLS |
| 2ba-C2 | + | - | + | - | BLS |
| 2a-D | + | - | + | - | BLS |
| 2a-E | + | - | + | - | BLS |
| 2a-F | + | - | + | - | BLS |
| other *P. syringae* strains |  |  |  |  |  |
| Clade 2b | + | - | - | - | nd or clade 2b |
| Clade 2d | + | - | - | + | clade 2d |

<sup>1</sup> Phylogroups, clades and clusters are named according to Berge et al 2014, Newberry et al 2019, Lacault et al 2020, and Djitro et al 2022a.
<sup>2</sup> qPCR as described in Lacault et al 2024.
<sup>3</sup> (VCZ) strain having a narrow host range of cucurbits and no disease on adult plant, (BLS) strain having a wide host range of cucurbits causing BLS symptoms on adult plants, according to Lacault et al 2020 and 2024. (nd) not determined.

Several tools are available to characterize this diversity and to detect these strains in zucchini seed production. A multilocus sequence analysis (MLSA) scheme, based on seven housekeeping genes (*gapA*, *gltA*, *gyrB*, *rpoB, Psyr3420*, *Psyr4880*, and *Psyr3208*) resolves the strains into the clades described above (Lacault et al. 2020; 2024). For diagnostic purposes, a qPCR test targeting *sylC* detects all strains that produce syringolin A, including VCZ and BLS strains as well as some clade 2b and 2d strains (Amrein et al., 2004; Manceau et al. 2011; Lacault et al. 2024). A multiplex qPCR test combining *sylC* with *avrRpt2*, *hopZ5*, and a clade-2d phylogenetic marker gene (MCA5971736.1) further identifies the host range of infecting strains (Lacault et al. 2024). Furthermore, a dual-probes qPCR test based on *sylC,* distinguishes clades 2a and 2ba, which include all the strains responsible for the most severe seedling symptoms (Willmann et al. 2026).

Because chemical control options of bacterial diseases in the field remained limited and ineffective in the long term (Miller et al 2022), seed health is central for preventing epidemics (Gitaitis and Walcott 2007). Seed compagnies therefore rely primarily on testing to keep contaminated lots off the market, most often using a bioPCR protocol (Schaad et al. 1995) that enriches bacteria from seed extracts before *sylC* qPCR detection (Manceau et al. 2011). A validated grow-out test, with isolation of bacteria and identification by pathogenicity test is also available for *Cucurbita pepo* seeds (Lybeert et al. 2022). Contaminated lots can be disinfected with chemical or hot water treatments (Lybeert et al. 2022), but this approach has its limits. First, many seed lots produced worldwide are infected, and this trend has been on the rise over time. This situation therefore results in large volumes of material that must be disinfected, which is time-consuming and costly. Furthermore, some seed lots remain infected even after several successive disinfection treatments. Preventing infection in seed crops would avoid these costs, which makes understanding how VCZ and BLS strains are transmitted to zucchini seeds an essential step toward more effective preventive measures.

The transmission of bacteria to seeds has been described for few pathosystems. Bacterial pathogens can colonize seeds through three routes: *via* the vascular system, via the flower through the stigma and pistil, and through contact with the fruit (Maude 1996). These three pathways have been reported for *Xanthomonas citri* pv. *fuscans* on beans (Chen et al. 2021). In the case of a pathosystem more closely related to zucchini, such as watermelon seeds infected by *Paracidovorax citrulli* (Bergmann et al. 2026), transmission to the seeds can occur through the floral route at the time of pollination and by contact via the pericarp up to two weeks after anthesis (Dutta et al. 2012; 2015). Fruits whose seeds have been contaminated through the flowers usually remain asymptomatic (Walcott et al. 2003). The bacteria are then confined to the internal tissues of the seed, which allows them to survive longer and resist disinfection treatments. On the contrary, infections via the pericarp are accompanied by symptoms. While there is no difference in the levels of seed infection compared to the floral route, the bacteria that colonize the seeds through the pericarp are localized in the outer tissues and are less able to survive over time and during seed treatments (Dutta et al. 2016). To date, there have been no reports on how *P. syringae* is transmitted to zucchini seeds.

The objectives of this study were to provide insights into the epidemiology of these diseases, VCZ and BLS. We addressed these questions: (i) which types of disease (VCZ or BLS) currently predominate in zucchini seed lots? Have BLS strains, become dominant compared to the historical strains? (ii) what are the sources of inoculum for these strains in seed crops? (iii) what are the routes of transmission, and are they the same for the VCZ and BLS strains; and (iv) does the infection process from seed to seedling follow the same dynamics depending on the type of strain? To answer these questions, we used survey-based and experimental approaches.

## Materials and methods

### Bacterial strains

A set of 37 strains were isolated during this work from surveyed fields: 19 strains from zucchini leaves and pollen, 13 strains from weeds and five strains from the produced hybrid zucchini seed lots (Supplementary Table S1). P99 and P66 strains were selected from previous work to represent VCZ and BLS strains, respectively (Lacault et al. 2020). To trace the bacterial strains used in seed transmission experiments, spontaneous rifamycin resistant variants were selected by spreading bacterial suspensions at 1×10^9^ CFU/ml on plates of 10% TSA medium (1.7 g/liter tryptone, 0.3 g/liter soybean peptone, 0.25 g/liter glucose, 0.5 g/liter NaCl, 0.5 g/liter K_2_HPO_4_, and 15 g/liter agar) containing 200 µg/liter of rifamycin. After 48 h of incubation at 28°C single colonies were purified and checked for growth and pathogenicity. Strains P99 Rif^R^ and P66 Rif^R^ were selected among several variants to have the same behavior as the parental strains P99 and P66 (Supplementary Figure S1).

Bacterial strains were stored at -80°C in 40% glycerol in sterile deionized water. Strains were routinely cultured at 28°C on 10% TSA medium. Strain isolation was performed on LBCAL medium (2 g/liter of yeast extract, 5 g/liter of Bacto Peptone, 50 g/liter of saccharose, 1.5 g/liter of boric acid, 15 g/liter of agar and 2 ml of NaOH 1 N, 40 mg/liter of cephalexin, 50 mg/liter of lincomycin and 50 mg/liter of cycloheximide). Bacterial suspensions were prepared in sterile deionized water from cultures grown on 10% TSA for 24 h at 28°C. Suspensions were adjusted to an optical density (OD) of 0.1 at a wavelength of 650 nm, corresponding to 1×10^8^ CFU/ml. Suspensions were diluted to the appropriate concentration. Quantification of bacterial population size was performed by dilution and plating on 10% TSA. The number of colonies was counted after 48 h of incubation at 28°C. P99 Rif^R^ and P66 Rif^R^ were grown on 10% TSA or LBCAL supplemented with rifamycin at 50 mg/liter.

### qPCR tests used to detect and identify *P. syringae* strains

A qPCR test targeting a fragment of a syringolin biosynthetic gene, *sylC,* was used to detect all VCZ and BLS strains and other *P. syringae* strains belonging to clades 2b and 2d (Table 1; Lacault et al 2024). A multiplex qPCR test (Lacault et al 2024), combining *sylC* test with three qPCR tests based on two genes encoding type III effectors (*avrRpt2* and *hopZ5*) and a marker of clade 2d encoding a chemotaxis methyl-accepting receptor, was used to identify the strains. All these qPCR tests were based on a Taqman technology with probes labelled with different reporter and quencher fluorophores (Supplementary Table S2). Positive qPCR results for *sylC* and *avrRpt2* correspond to VCZ strain, while positive qPCR results for *sylC* and *hopZ5* correspond to BLS strain. Positive qPCR results for *sylC* and clade 2d marker indicate a clade 2d strain (Lacault et al 2024). The multiplex qPCR test was performed in a final volume of 10 µl containing 5 µl of Sso Advanced Universal Probes Supermix (Bio-Rad, Marne-la-Coquette, France), 600 nM of each primer, 200 nM of each probe and 1 µl of DNA. The amplification program was 3 min at 95°C, followed by 40 cycles of 15 s at 95°C and 30 s at 60°C. The qPCR test for *sylC* alone was also performed in a final volume of 20 µl containing 10 µl of MasterMix buffer (Eurogentec, Seraing, Belgium), 900 nM of each primer, 250 nM of probe, and 5 µl of DNA with an amplification program of 10 min at 95°C followed by 40 cycles of 15 s at 95°C and 15 s at 60°C. All qPCR tests were performed in a Bio-Rad CFX96 Touch thermocycler and analyzed with Bio-Rad CFX Manager 3.1 software. For each reaction, the standard curve, the efficiency of the qPCR test (E= 10^(−1/slope)^), and the correlation coefficient were determined using boiled bacterial dilutions (10^8^ to 10^3^ CFU/ml) of strains P99, P66 and P129 for the qPCR tests for *sylC* and *avrRpt2*, for *hopZ5,* and for clade-2d strains, respectively.

### Seed testing

Zucchini seed samples were soaked in sterile phosphate-buffered saline (PBS, Sigma-Aldrich, Schnelldorf, Germany) supplemented with 100 µl/liter of Tween20 for 2 h at constant agitation (105 rpm) at room temperature at a ratio of 3 ml per g of seed or 1 ml per individual seed. For direct qPCR testing, DNA was extracted directly with QuickPick SML Kit (Bio-Nobile, Turku, Finland) from one ml of seed extract after centrifugation (15,000 G for 20 min), according to the supplier’s instruction in an automated system (Caliper Zephyr, PerkinElmer, Villebon-sur-Yvette, France). The DNA was then eluted from the magnetic beads using 30 µl of elution buffer. For each sample, 5-µl DNA aliquots were tested with the *sylC* qPCR test and 1-µl DNA aliquots were tested with the multiplex qPCR test to identify the type of strains. The detection threshold (minimal quantity detected in a sample) of the multiplex qPCR test was 1 × 10^5^ and 5 × 10^5^ CFU per subsample of 100- and 500-seeds, respectively and that of the *sylC* qPCR test was 50 times lower than the multiplexed reaction due to a tested volume five time bigger and a detection threshold 10 times lower in simplex than in multiplex (Lacault et al. 2024). For bioPCR testing, one ml of seed extract was plated on LBCAL medium. After 24-h growth, all bacterial cells were collected in 3 ml of 0.85% PBS. One ml of this suspension was centrifuged at 13,000 g for 5 min. The pellet was lysed with 500 µl of 0.5 N NaOH, incubated at 65°C for 10 min and then placed on ice. The DNA solution was diluted to the hundredth in 20 mM Tris-HCl buffer pH 8 and analyzed directly by qPCR. Each sample was tested one to three times independently using the *sylC* qPCR test. The detection threshold of a bioPCR test was 2.5 × 10^2^ and 1.25 × 10^3^ CFU per subsample of 100- and 500-seeds, respectively. For dilution-plating, used for experimental transmission of rifamycin resistant strains, 500 µl of crude extract and 50 µl of serial dilutions were plated on 10% TSA supplemented with rifamycin. Typical P99 Rif^R^ or P66 Rif^R^ colonies were numbered after 48-h growth at 28°C. The detection threshold of dilution plating varied from 2 to 22 CFU per subsample of 1- to 50-seeds, respectively.

### Calculation of the infection rate of a seed lot

The infection rate of seed lots was determined by analysis of multiple subsamples. The calculation was made either (i) from the number of infected subsamples for degressive samples using most probable number tables (Swaroop 1951) (ii) or by a probability calculation based on the number of healthy groups among (N) groups of identical size each containing (n) seeds (Maury et al. 1986).

### Characterization of infections present in seed lots produced in different countries

The multiplex qPCR test was used to characterize the *P. syringae* strains present in the zucchini seed production chain between 2018 and 2020. We tested 27 DNAs sent by industry. These DNAs corresponded to enriched seed extracts that had tested positive with the *sylC* bioPCR test during seed testing.

### Zucchini seed crop survey

In 2018 and 2019, the same parental seed lots were sown to produce hybrid seeds in two different regions (i) Limagne (<u>45° 57′ 33″ North,</u> <u>3° 35′ 38″ East</u>), and (ii) the Rhône valley (<u>45° 00′ North, 4° 50′ East</u>). Parental and hybrid seed lots (1,000 to 2,000 seeds per seed lot with subsamples of 100 to 500 seeds) were tested for bacterial infection. Seed crops were inspected for the presence of symptomatic plants. Male parents were sown 15 days before female parents. Plot sampling was carried out one month after sowing the females. Plots were covered in several diagonal lines. When a putative symptom was observed, the symptomatic plantlet or plant fragment was sampled to check bacterial infection. Female flowers of female parent and male flowers of male parent were also sampled, as well as fragments (or total aerial part for small specimens) of weeds and volunteers, symptomatic or not.

### Plant sample testing

Pollen was collected from male flowers and pistil from female flowers, a fragment of asymptomatic tissue located at the edge of the symptom was cut from symptomatic samples and a fragment of seedling or leaf from symptomless samples. For all samples, a fragment of equivalent size (approximately 3 × 6 cm) was ground using a mixing paddle (MixWell, Alliance Bio Expertise, France) with 5 ml of sterile water. A 200-µL aliquot of each extract was stored in 40% glycerol at -80°C and 1 ml of the extract was centrifuged at 15,000 g for 10 min. DNA was extracted from the pellet using an extraction kit (QuickPick SML Plant DNA, Bio-Nobile, Finland). After cell lysis, up to 96 samples could be processed simultaneously using an automated system (Caliper Zephyr, PerkinElmer). DNA was suspended in 30 µl of deionized water. Five µl of each sample were tested in duplicate using the *sylC* qPCR test in a total volume of 20 µL. Bacterial population sizes were determined for each sample. The detection threshold for qPCR was 150 bacteria per leaf fragment. For positive samples, the macerate stored at -80°C was plated on LBCAL medium and incubated at 28°C for 24 h to isolate strains.

### Phylogenetic placement of new isolates

Thirty-seven strains isolated from field survey and harvest were typed with three genes (Psyr3420, Psyr4880 and Psyr3208) as described in Lacault et al. 2024. Primer pairs Psyr3208-F and Psyr3208-R, Psyr3420-F and Psyr3420-R, and Psyr4880-F and Psyr4880-R (Supplementary Table S2) amplified of 883-, 343-, and 416- bp fragments, respectively. PCR tests were performed in a 50-µL volume containing 10 µl of buffer, 0.4 U/µl of Taq polymerase (GoTaq, Promega, Charbonnières-les-Bains, France), 200 µM of dNTPs, 0.5 µM of each primer, and 5 µl of boiled bacterial suspension. For *Psyr3208* primers, amplification conditions were 5 min at 94°C followed by 20 cycles of 30 s at 94°C, 30 s at 60°C (with a 0.5°C decrease at each cycle), and 1 min at 72°C, then 15 cycles of 30 s at 94°C, 30 s at 50°C, and 1 min at 72°C. For *Psyr3420* or *Psyr4880* primers, PCR conditions were 5 min at 94°C followed by 35 cycles of 30 s at 94°C, 30 s at 62°C (for *Psyr3420*) or 66°C (for *Psyr4880*), and 1 min at 72°C. A final step of 10 min at 72°C was added to achieve elongation for all PCR tests. Amplified fragments were Sanger sequenced with the forward and reverse primers, and with MF- and MR-primers for *Psyr3208* (Genoscreen, Lille, France). Additional sequencing primers, Psyr3208-MF and Psyr3208-MR (Supplementary Table S2), designed in the middle of the *Psyr3208* fragment, were used with Psyr3208-R and Psyr3208- F, respectively, in order to increase the length of double-stranded sequences. Geneious version 9.1.7 software was used to assemble, align, orientate, and trim sequences according to the reading frame. Sequences of partial coding DNA sequences (CDSs) were deposited in GenBank under the accession numbers PV754503 to PV754602, and were 825- (*Psyr3208*), 318- (*Psyr3420*), and 309-bp (*Psyr4880*) long. Phylogenetic analyses were performed on the concatenated sequence (1452 bp) of the three genes for 37 strains newly isolated during the survey (Supplementary Table S1). The tree was constructed by including the concatenated sequences of strains from previous studies (Lacault et al 2020; 2024; Newberry et al. 2019), and of strains representing the diversity of the *P. syringae* species complex (Supplementary Table S3). Sequences from the *P. viridiflava* strain (ICMP 3272) were used to infer a maximum likelihood (ML) tree using the Tamura-Nei model with gamma distribution of invariant sites (G + I) and 1000 bootstraps (MEGA 7, Kumar et al., 2016).

### Transmission experiments of P99 Rif^R^ and P66 Rif^R^ strains to zucchini seeds

A commercial zucchini seed lot of the cultivar Divonne was used in the seed transmission tests for P99 Rif^R^ and P66 Rif^R^ strains. This lot was free from VCZ and BLS infections based on the negative analysis of 2,000 seeds with the qPCR test targeting *sylC*. Seeds were sown in individual 9 × 9 cm pots containing substrate 4 (Klasmann-Deilmann, Ruy Montceau, France). Three weeks after sowing, seedlings were transplanted into 15 L pots. Plants were watered daily using a fertilizer irrigation system with an NPK ratio of 12/7/25 and an effective cation exchange value of 1.8. At anthesis, one just opened female flower per plant was cross-pollinated early in the morning with pollen from another plant. Only one flower was fertilized per plant. Three infection routes were tested using water (negative control) or calibrated suspensions (1 × 10^7^ CFU/ml) of P99 Rif^R^ and P66 Rif^R^. Two independent experiments (I and II) were conducted, each with at least five plants per treatment. To limit the cases of abortion observed in the floral and vascular pathways, suspensions at 1 × 10^5^ CFU/ml were inoculated in a third experiment (III) for both pathways. For the vascular route, the female flower stalk was infiltrated twice with a needle (Terumo 18 g pink 1.2 mm × 38 mm) previously filled with bacterial suspension by capillary action. The two injections were made perpendicularly to optimize the number of vessels inoculated. The infiltrated volume was estimated to be 2 µL. For the floral route, a 50-µL drop of bacterial suspension was deposited on the stigma of the female flower just after pollination. For the contact route, a 5 × 5 cm square of sterile filter paper soaked with 200 µl of bacterial suspension was applied to the pericarp of a one-week-old fruit. Plants were cultivated until fruit ripening, i.e. 75 ± 5 days, according to greenhouse zucchini standards. At harvest, the fruit surface was disinfected with 70% alcohol. For each fruit, only mature seeds were extracted, rinsed with sterile distilled water and dried on filter paper at room temperature for one week. All seeds from a fruit constituted a seed lot (up to 300 seeds per fruit). Seed lots were stored at 9°C until analysis. Detection of P99 Rif^R^ and P66 Rif^R^ was performed by dilution plating on selective TSA 10% medium (supplemented with Rifamycin 50 mg/liter). The detection threshold was 1 CFU/ml when plating 1 ml of crude extract. For each seed lot, a subsample of 50 seeds was analyzed. In case of a positive sample, five subsamples of 10 seeds and five single seeds were analyzed and the infection rate of the lot was estimated using the tables of Swaroop (1951). In case of a negative subsample, two more subsamples of 50 seeds were analyzed and the infection rate of the lots was then estimated using Maury’s formula with N = 3 and n = 50. For small lots, analyses were performed directly on subsamples of 10 seeds or single seeds and the infection rate was determined using Maury’s formula.

### Experiments on the transmission of VCZ and BLS strains from seed to seedling

Bacterial transmission to seedlings was studied by comparing population sizes distribution in seeds and seedlings using three highly infected seed lots (X, Y, and Z). X was a mix of infected zucchini seeds, Y corresponded to pumpkin seeds infected with a VCZ strain, and Z was a zucchini seed lot inoculated by the vascular pathway with strain P66 Rif^R^ (BLS strain). Because seed testing is a destructive method, 30-seed subsamples per seed lot were individually analyzed as previously described or sown in individual 9 × 9 cm pots containing substrate 4 in a mini greenhouse at saturated relative humidity. For lot X, the experiment was repeated three times. After seven days, seedlings at the cotyledonary stage were collected by cutting them individually at the base of the collar. Plant samples were placed individually in a plastic bag containing 5 ml of sterilized water and ground for 5 min. For each sample, 1 ml of the plant extract and 50 µl of appropriate dilutions were plated on LBCAL for X and Y seed lots or on medium supplemented with rifamycin for the Z seed lot. Plates were incubated for 24 h at 28°C. Typical *P. syringae* colonies were counted and their identity was verified using *sylC*-qPCR test.

### Statistical tests

Comparisons of proportions of infected seeds and seedlings were based on a Pearson’s χ² test run on biostaTGV (https://biostatgv.sentiweb.fr/).

## Results

### *Cucurbita pepo* seed lots infected with *Pseudomonas syringae* strains in different countries overwhelmingly contained VCZ strains

DNA samples from seed lots that tested positive for the presence of *P. syringae* strains with the *sylC* marker were characterized using a multiplex test (Lacault et al., 2024) to determine their identity (VCZ or BLS types or to clade 2d) (Table 2). More than 66% (18 out of 27) of the seed lots were exclusively composed of VCZ strains with narrow host range (*sylC* + *avrRpt2*). Three samples contained exclusively BLS strains (*sylC* + *hopZ5*). Four seed lots contained DNA from both types of strains. Finally, two seed lots were infected by strains belonging to clade 2d.

**Table 2.** Identification of strain types present in infected *Cucurbita* seed lots produced in different countries between 2018 and 2020.

| Sample number | Species | Type | Country | Year | bioPCR<sup>1</sup> |  | multiplex qPCR<sup>2</sup> |  |  | type of VCZ<sup>3</sup> |
| --- | --- | --- | --- | --- | --- | --- | --- | --- | --- | --- |
|  |  |  |  |  | *sylC* | *sylC* | *avrRpt2* | *hopZ5* | PG 2d |  |
| Lot 1 | *C. pepo* subsp. *pepo* | zucchini | China | 2018 | + | + | + | - | - | VCZ |
| Lot 2 | *C. pepo* subsp. *pepo* | pumpkin | China | 2018 | + | + | + | - | - | VCZ |
| Lot 3 | *C. pepo* subsp. *pepo* | pumpkin | China | 2018 | + | + | + | - | - | VCZ |
| Lot 4 | *C. pepo* subsp. *pepo* | pumpkin | China | 2018 | + | + | + | - | - | VCZ |
| Lot 5 | *C. pepo* subsp. *pepo* | pumpkin | China | 2018 | + | + | + | - | - | VCZ |
| Lot 6 | *C. pepo* subsp. *pepo* | pumpkin | China | 2018 | + | + | + | - | - | VCZ |
| Lot 7 | *C. pepo* subsp. *pepo* | pumpkin | China | 2018 | + | + | + | - | - | VCZ |
| Lot 8 | *Cucurbita* sp. | unknown | unknown | 2018 | + | + | + | - | - | VCZ |
| Lot 9 | *C. pepo* subsp. *pepo* | zucchini | China | 2018 | + | + | + | - | - | VCZ |
| Lot 10 | *C. pepo* subsp. *pepo* | zucchini | France | 2018 | + | + | + | - | - | VCZ |
| Lot 11 | *C. pepo* subsp. *pepo* | pumpkin | China | 2018 | + | + | + | - | - | VCZ |
| Lot 12 | *C. pepo* subsp. *pepo* | pumpkin | China | 2018 | + | + | + | - | - | VCZ |
| Lot 13 | *C. pepo* subsp. *pepo* | pumpkin | China | 2018 | + | + | + | - | - | VCZ |
| Lot 14 | *C. pepo* subsp. *pepo* | pumpkin | China | 2018 | + | + | + | - | - | VCZ |
| Lot 15 | *C. pepo* subsp. *pepo* | zucchini | China | 2018 | + | + | + | - | - | VCZ |
| Lot 16 | *C. pepo* subsp. *pepo* | zucchini | China | 2018 | + | + | + | - | - | VCZ |
| Lot 17 | *C. moschata* | butternut | Inde | 2018 | + | + | + | - | - | VCZ |
| Lot 18 | *C. pepo* subsp. *pepo* | pumpkin | China | 2018 | + | + | + | - | - | VCZ |
| Lot 19 | *C. pepo* subsp. *pepo* | zucchini | China | 2018 | + | + | + | + | - | VCZ + BLS |
| Lot 20 | *C. pepo* subsp. *pepo* | zucchini | China | 2018 | + | + | + | + | - | VCZ + BLS |
| Lot 21 | *C. pepo* subsp. *pepo* | zucchini | France | 2018 | + | + | - | - | + | clade 2d |
| Lot 22 | *C. pepo* subsp. *pepo* | zucchini | France | 2018 | + | + | - | - | + | clade 2d |
| Lot 23 | *C. pepo* subsp. *pepo* | zucchini | China | 2018 | + | + | - | + | - | BLS |
| Lot 24 | *C. moschata* | butternut | Thailand | 2018 | + | + | - | + | - | BLS |
| Lot 25 | *C. pepo* subsp. *pepo* | zucchini | Chile | 2018 | + | + | - | + | - | BLS |
| Lot 26 | *C. pepo* subsp. *pepo* | zucchini | China | 2020 | + | + | + | + | - | VCZ + BLS |
| Lot 27 | *C. pepo* subsp. *pepo* | zucchini | China | 2020 | + | + | + | + | - | VCZ + BLS |
| number of positive samples |  |  |  |  | 27 | 27 | 22 | 7 | 2 |  |

<sup>1</sup> Samples were analyzed by bioPCR: culture-enriched seed extracts were tested in duplicate (2 x 5 µl) using the *sy/C* qPCR test.
<sup>2</sup> Same culture-enriched seed extracts were analyzed in triplicate (3 x 1 µl) using the multiplex qPCR test.
(+) indicated a Ct value was obtained for at least 2 repeats of each qPCR target and (-) indicated that no Ct value was obtained for each qPCR target.
<sup>3</sup> (VCZ) strain having a narrow host range of cucurbits and no disease on adult plant, (BLS) strain having a wide host range of cucurbits causing BLS symptoms on adult plants.

### The study of inoculum sources of *P. syringae* strains in seed production revealed that parental lots were infected with low levels of non-culturable cells of *P. syringae* from PG2. Some of these infections were identified as BLS strains but no VCZ infections were evidenced

The origin of the strains detected in the seed lots was first assessed by analyzing (i) the parental lines used to generate these hybrids. None of the parental line samples tested positive with the culture-based test (*sylC* bio-qPCR test) (Table 3). However, positives samples were detected with the *sylC* qPCR test in five out of six parental lines when DNA from these corresponding lots was used (Table 3 and Supplementary Table S4). Three of these parental lines were infected with BLS strains, while infections of two parental lines could not be characterized, either because bacterial population sizes were too low to be detected using the multiplex qPCR test or because infections corresponded to clade-2b strains (Supplementary Table S4). It was not possible to conclusively determine whether some parent lots were contaminated with VCZ strains. The infection rates of the parental seed lots ranged from 0.08% to 0.22%, which corresponds to one infected seed among 1,250 seeds or one infected seed among 454 seeds, respectively.

**Table 3.**
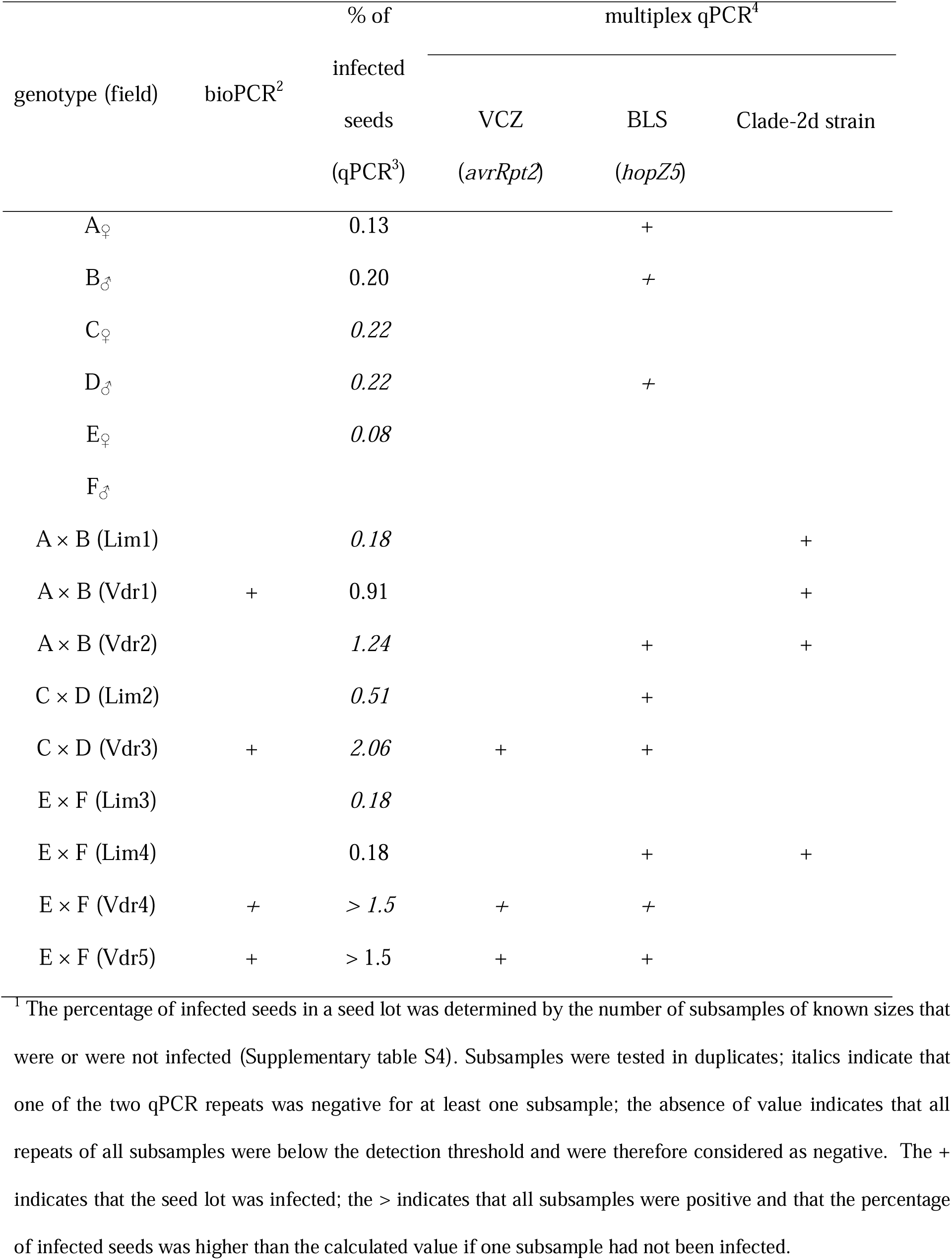
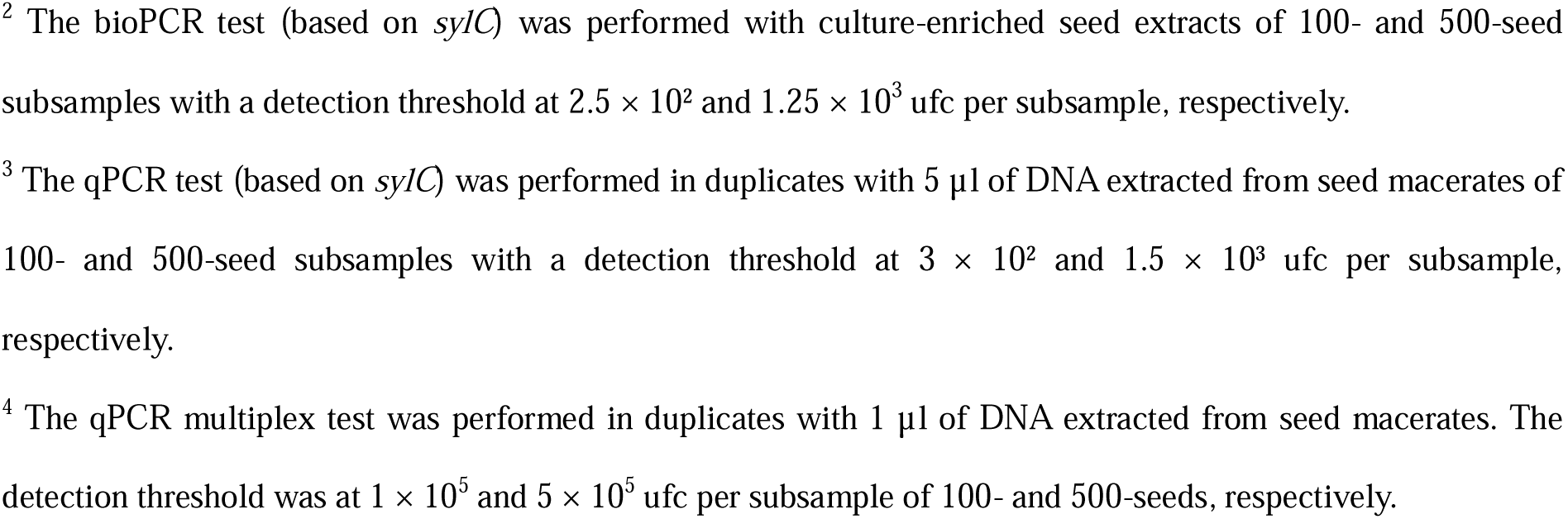
Percentage^1^ of seeds infected with *Pseudomonas syringae* strains in parental and hybrid seed lots.

| genotype (field) | bioPCR<sup>2</sup> | % of infected seeds (qPCR<sup>3</sup>) | multiplex qPCR<sup>4</sup> |  |  |
| --- | --- | --- | --- | --- | --- |
|  |  |  | VCZ (*avrRpt2*) | BLS (*hopZ5*) | Clade-2d strain |
| A<sub>♀</sub> |  | 0.13 |  | + |  |
| B<sub>♂</sub> |  | 0.20 |  | + |  |
| C<sub>♀</sub> |  | 0.22 |  |  |  |
| D<sub>♂</sub> |  | 0.22 |  | + |  |
| E<sub>♀</sub> |  | 0.08 |  |  |  |
| F<sub>♂</sub> |  |  |  |  |  |
| A × B (Lim1) |  | 0.18 |  |  | + |
| A × B (Vdr1) | + | 0.91 |  |  | + |
| A × B (Vdr2) |  | 1.24 |  | + | + |
| C × D (Lim2) |  | 0.51 |  | + |  |
| C × D (Vdr3) | + | 2.06 | + | + |  |
| E × F (Lim3) |  | 0.18 |  |  |  |
| E × F (Lim4) |  | 0.18 |  | + | + |
| E × F (Vdr4) | + | > 1.5 | + | + |  |
| E × F (Vdr5) | + | > 1.5 | + | + |  |

<sup>1</sup> The percentage of infected seeds in a seed lot was determined by the number of subsamples of known sizes that were or were not infected (Supplementary table S4). Subsamples were tested in duplicates; italics indicate that one of the two qPCR repeats was negative for at least one subsample; the absence of value indicates that all repeats of all subsamples were below the detection threshold and were therefore considered as negative. The + indicates that the seed lot was infected; the > indicates that all subsamples were positive and that the percentage of infected seeds was higher than the calculated value if one subsample had not been infected.
<sup>2</sup> The bioPCR test (based on *sy/C*) was performed with culture-enriched seed extracts of 100- and 500-seed subsamples with a detection threshold at $2.5 \times 10^2$ and $1.25 \times 10^3$ ufc per subsample, respectively.
<sup>3</sup> The qPCR test (based on *sy/C*) was performed in duplicates with 5 μl of DNA extracted from seed macerates of 100- and 500-seed subsamples with a detection threshold at $3 \times 10^2$ and $1.5 \times 10^3$ ufc per subsample, respectively.
<sup>4</sup> The qPCR multiplex test was performed in duplicates with 1 μl of DNA extracted from seed macerates. The detection threshold was at $1 \times 10^5$ and $5 \times 10^5$ ufc per subsample of 100- and 500-seeds, respectively.

### The *P. syringae* strains isolated in plots from parental zucchini plants and other surrounding weeds belonged to clades 2b and 2d, none were VCZ nor BLS strains

The seed crops (open fields) were then monitored and no outbreaks of disease were observed. The overall appearance of the crops was healthy. We surveyed the plots along several diagonals to look for anything that could resemble symptoms such as necrotic spots on the leaves or growth delay. Among the 582 samples collected during the field survey, 74 were colonized by *P. syringae* bacteria (Table 4) with population sizes ranging from 50 to 5 × 10^7^ CFU/sample (Supplementary table S5). Thirty-two strains were isolated from the samples and MLSA results showed they belong to clades 2b and 2d (Figure 1). These strains were phylogenetically diverse but some of them clustered together. One cluster of clade-2b strains (containing strains 108, 110, 122, 107, and 106) was composed of strains harboring the same sequence type and all were isolated from the same field and mainly from weeds and volunteers suggesting an environmental origin (Figure 1 and Supplementary table S5). Two clade-2d clusters (the first cluster containing strains P134, 428A, 433C, 410, 408, 133, and 53 and the second cluster containing strains 439A, 440B, 438A, 436A, 143I, 142I, and 32) each corresponding to a unique sequence, included strains isolated from different plots, mainly from zucchini plants belonging to different parental lines, and some were recovered from pollen of one male parent. Furthermore, one of these clusters contained one strain previously isolated from zucchini seed lots (P134). Although some evidence would appear to support the hypothesis of a seed origin, the inoculum sources of these clusters, within clade-2d, remain unidentified, as they do for all the other phylogenetically diverse clade-2b and -2d strains. Despite parental seed lots A, B, and D carried infection remnants of BLS strains, we did not isolate any BLS strains from the 168 zucchini symptomatic samples of these three parental lines. No VCZ strains were recovered from field samples.

**Figure 1.**
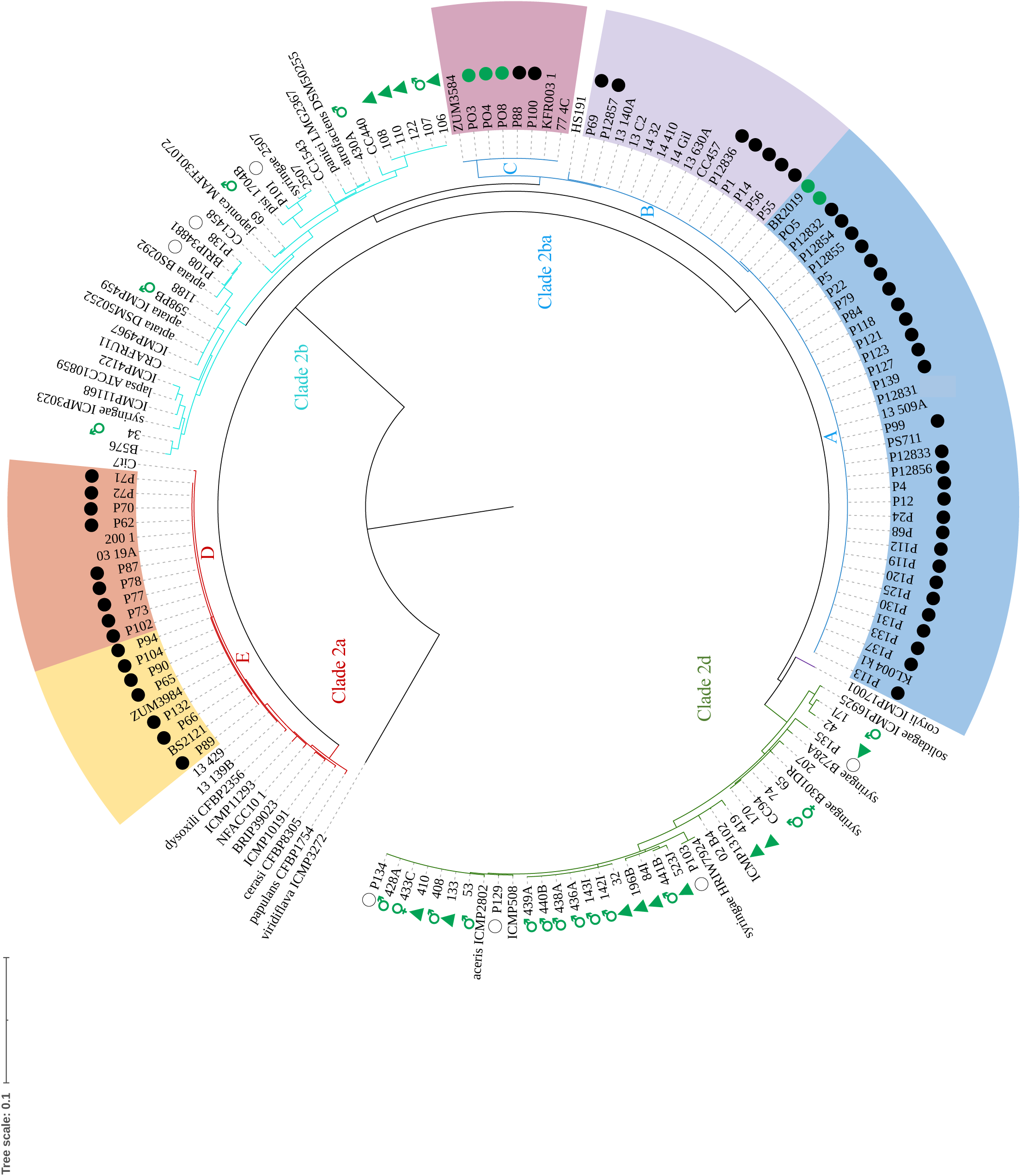
Phylogenetic relationships of the *Pseudomonas syringae* strains isolated from field survey. The maximum likelihood (ML) tree was based on concatenated partial sequences of Psyr3208, Psyr3420, and Psyr4880 (1,452 bp) of 36 strains isolated from the survey (Supplementary Table S1) and a reference collection of 120 strains (Supplementary Table S2). Clades are indicated based on several studies (Berge et al. 2014; Bull and Koike 2015; Newberry et al. 2019), and clusters according to Lacault et al. (2020). Green symbols are used for strains isolated during the survey and black is used for our reference isolate collection. Solid circles indicate VCZ strains and empty circles are used for other *P. syringae* strains. Male and female symbols indicate isolation from male or female zucchini plants, and a triangle indicates weeds or volunteers.

**Table 4.** Positive samples (*sylC* qPCR test) from the field survey.

| genotype | plot | Number of samples (infected/total) |  |  |
| --- | --- | --- | --- | --- |
|  |  | ♀ zucchini | ♂ zucchini | weed |
| A × B | Lim1 | 3/18 | 13/16 | 3/13 |
| A × B | Vdr1 | 0/30 | 0/13 | 1/18 |
| A × B | Vdr2 | 2/34 | 2/13 | 2/33 |
| C × D | Lim2 | 0/18 | 0/28 | 0/13 |
| C × D | Vdr3 | 0/20 | 0/16 | 0/20 |
| E × F | Lim3 | 0/6 | 0/20 | 3/15 |
| E × F | Lim4 | 0/12 | 1/16 | 0/20 |
| E × F | Vdr4 | 3/20 | 7/23 | 15/56 |
| E × F | Vdr5 | 4/28 | 4/30 | 11/33 |

### More than 66% of the hybrid seed lots produced in the Rhône Valley and in Limagne were infected with BLS strains and 33% with VCZ strains. Contamination rates were higher in plots located in Rhône Valley than in those located in Limagne

The percentage of infected seeds was higher in plots located in the Rhône Valley than in Limagne indicating that conditions in the Rhône Valley were more favorable to infection of seeds by VCZ strains (Table 3, Supplementary Table S4). VCZ strains were recovered only in seed lots produced in this region. The origin of infection by VCZ strains remains uncertain, as all hybrid seed lots infected with this type of strain originate from parental lines that were negative to VCZ test. In the case of A × B hybrid seed lots, despite they derived from parental seed lots both infected with BLS strains, BLS infection was detected in the seeds of only one of the three fields. The C × D hybrid seed lots, derived from a male parent whose seeds showed traces of infection with a BLS strain, were infected with a strain of the same type in both production regions, strongly suggesting that this infection could originate from the seeds. Most of the E × F hybrid seed lots harbored mixed infections that could not be traced. Five VCZ strains could be isolated from these infected hybrid seed lots. Lastly, clade-2d strains were detected in hybrid seed lots produced in both regions but no strains could be isolated to check if they corresponded to strains recovered from the environment of the corresponding field (i.e. clade- 2d strain clusters, Figure 1).

### Inoculation of zucchini flowers or fruits showed that VCZ and BLS strains can colonize seeds by vascular and floral pathways but that only the BLS strain transmitted to seeds by external contact with fruit

Similar results were obtained in two independent experiments (experiments I and II Figure 2; Supplementary Table S6) with an inoculum dose at 1×10^7^ CFU/ml although more fruits aborted in experiment II. The inoculated plants did not develop any foliar symptoms and fruits harvested at maturity were asymptomatic, except for the contact pathway with the BLS strain, where small, dry pustules developed beneath the inoculated patch on the pericarp (Supplementary Figure S2). In contrast, the aborted fruits showed symptoms and were heavily infected with the inoculated strains (data not shown). The contact route did not induce any abortions for both strains. For the floral and vascular pathways, a third experiment was conducted with an inoculum dose 100 times lower to reduce abortions. The lower inoculum dose decreased the number of infected seeds and reduced the number of abortions for the floral route and the vascular route for strain P99 Rif^R^ and strain P66 Rif^R^, respectively (Figure 2, experiment III; Supplementary Table S6). Globally, floral pathway was more successful to transmit the VCZ strain (P99 Rif^R^) to seeds with 10 infected seed lots out of 17 compared to 3/20 and 0/10 by the vascular and the external pathways, respectively. Furthermore, percentages of seeds infected by the VCZ strain were also higher for the floral route. In contrast, the BLS strain (P66 Rif^R^) transmitted through all three routes, with ratios of infected/tested seed lots of 5/16, 8/17, and 7/10 or floral, vascular, and pericarp routes, respectively (Figure 2).

**Figure 2.**
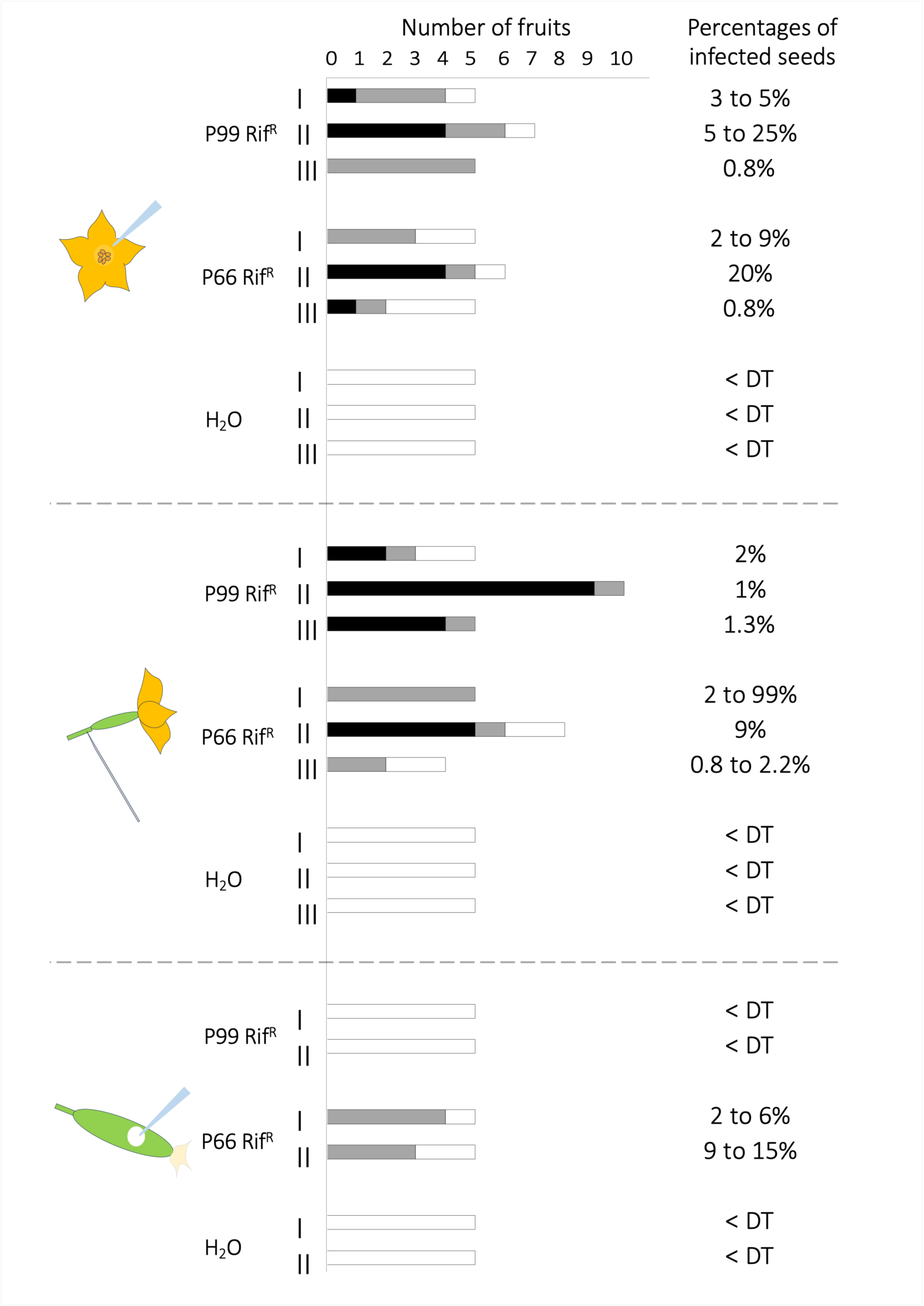
Experimental transmission of strains P99 Rif^R^ and P66 Rif^R^ to zucchini seeds by floral, vascular or contact pathways. Rifamycin resistant variants of a VCZ strain with a narrow cucurbit host range (P99 Rif^R^) and with a wide cucurbit host range (P66 Rif^R^) were inoculated by floral, vascular and contact pathway, using five plants per treatment and per experiment. Sterile distilled water was used as negative control. Two experiments (I and II) were performed using an inoculum calibrated at 1 × 10^7^ CFU/ml and a third experiment (III) using an inoculum calibrated at 1 × 10^5^ CFU/ml. Black color indicates abortion of the fruit, grey is for an infected seed lot, and white for no detected infection. Percentages of infected seed are given from the lower to the higher values.

### Comparison of the distribution of infection in seeds and seedlings showed that germination could reveal infections undetected in seeds and rule out low levels of infection that were not transmitted to the seedling

As analyses are destructive, seed lots were characterized by sampling between 30 and 30 × 3 seeds for direct testing and by similar sampling for testing after germination. The distribution of infection in the seeds was significantly different from that in the germinated seeds for the three seed lots X, Y, and Z with Pearson’s χ² test p-values 5.56 × 10^-9^, 1.18 × 10^-22^, and 4.99 × 10^-13^, respectively. Twenty-five and 20% of seed infections were not detected in lots X and Y, respectively, whereas in lot Z, 30% of infected seeds, possibly carrying bacterial low population sizes did not rise to infected seedlings (Figure 3). Major differences between the seed lots were the type of infection (VCZ for lots X and Y and BLS for lot Z) and the absence of any industrial process for lot Z, which was produced in our laboratory by vascular inoculation with strain P66 Rif^R^.

**Figure 3.**
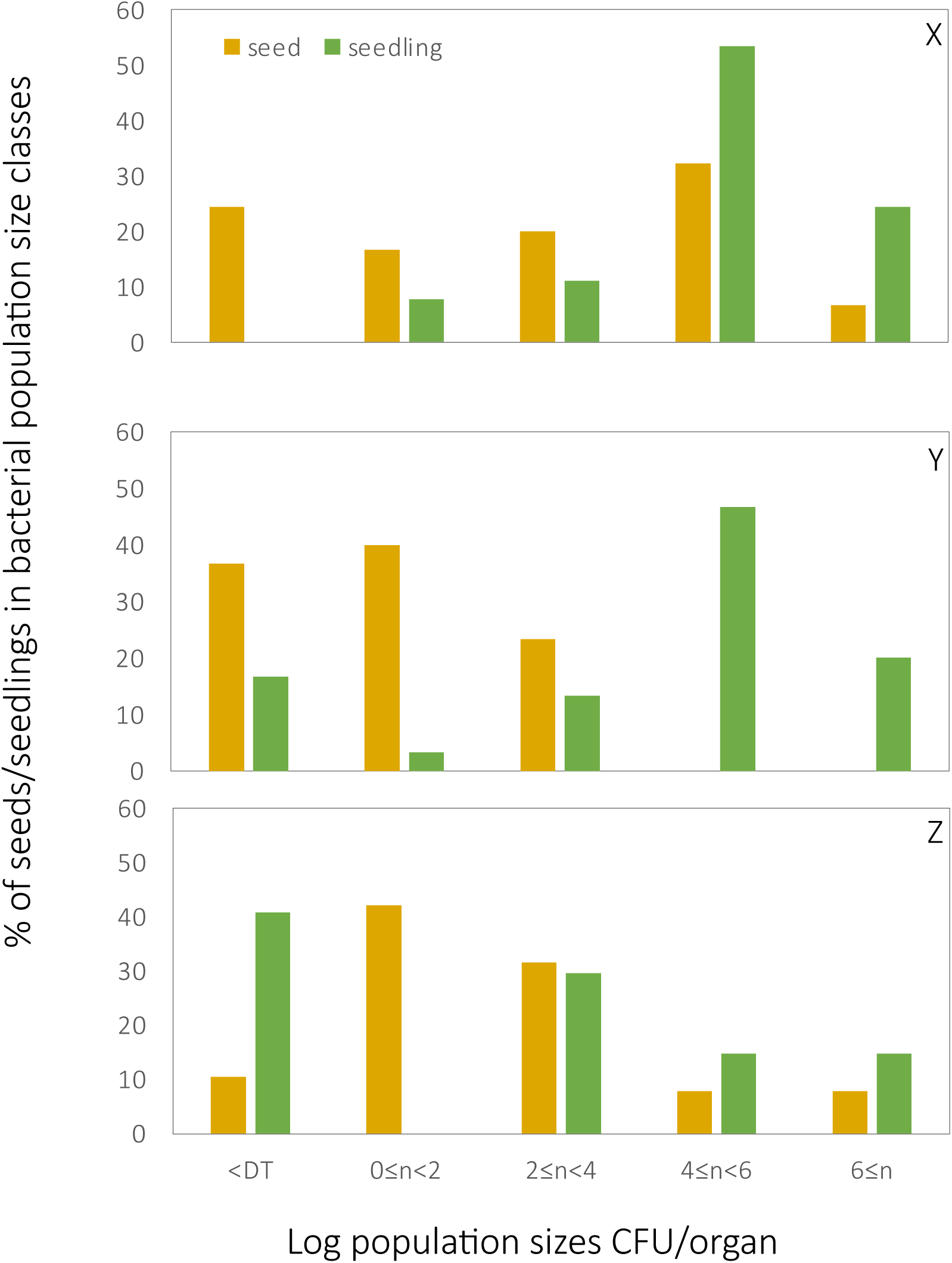
Seed testing with and without a germination step. Three seed lots (X, Y, and Z) were sampled for infection of individuals (seeds and 7-day old seedlings) before and after germination. A total of 90 individual seeds and 90 individual seedlings were tested for seed lot X, and 30 individual seeds and 30 individual seedlings for seed lots Y and Z. Samples were tested by dilution plating of seed or seedling extracts and subsequent colony identification using *sylC* qPCR. Detection threshold (DT) was two CFU for seed and five CFU for seedling.

## Discussion

This study aimed to improve our understanding of how zucchini seeds become infected by different strains of the *Pseudomonas syringae* species complex. Of particular interest were strains that cause symptoms exclusively on seedlings (VCZ) and those that cause symptoms on seedlings and leaf necrosis on adult plants (BLS). We used survey-based approaches on seed production plots and experimental approaches to transmit representative strains. The survey of seed crops did not allow us to reliably identify the sources of inoculum. Some findings nonetheless suggest the possibility of vertical transmission, while others highlight the importance of the environment in seed infection. With regard to vertical transmission, although no culturable bacteria were detected in any of the parental seed lots, *P. syringae* DNA was successfully extracted from seeds, showing that parental lineages were infected before being disinfected. We could therefore consider the possibility of residual infection by viable bacteria, as has been observed in some disinfected seed lots (Carisse et al. 2000; Temple et al. 2013). Some of these DNA contaminants were identified as BLS strains, as were the infections found in the hybrid seeds produced in both regions. Furthermore, the production region appeared to play an important role. Parallel trials conducted in the two regions consistently showed more frequent and higher levels of infection in the Rhône Valley than in Limagne, particularly for VCZ infections, which were recovered only from the seed lots produced in the Rhône Valley. Under our experimental conditions, the VCZ and BLS strains were able to infect the seeds through the vascular and floral pathways, and only the BLS strain was able to infect the seeds through the symptomatic pericarp. In a seed production context, the floral pathway could allow transmission *via* pollinators carrying infected pollen, and the vascular pathway could allow transmission through an infected female parent. Finally, individual seed and seedling testing by culture-based methods showed that seedlings infections exceeded seed infections by at least 20% in seed lots that had undergone an industrial process and were infected by VCZ strains. The opposite pattern was observed for a nontreated seed lot infected by a BLS strain, showing that some seedborne infections go undetected in seeds yet are still transmitted to seedlings.

This discrepancy raises the question of why undetected infections became apparent specifically in seedlings from the industrially processed lots contaminated with VCZ strains. In the untreated lot, some of the infections (possibly those with the lowest bacterial population sizes) were lost when the seeds germinated. This has been previously shown for bean seeds contaminated with low populations of *Xanthomonas citri* pv. *fuscans* that do not transmit to seedlings (Darrasse et al. 2007). Using a dilution-plating approach, the detection threshold in this experiment was better for the seeds than for the seedlings because the whole volume of the sample was plated for seeds whereas only one of the five ml used to grind seedlings was plated. Were there more infections in the seedlings than in the seeds from some lots because they had undergone an “industrial” process? Although industrial processes are confidential, it is known that they can include disinfection steps when the plant species and conditions warrant it. The ability of phytopathogenic bacteria to survive in seeds for decades (Bashan et al. 1982) suggests that they can establish resistant forms that escape eradication treatments (Leben 1974). *In vitro* studies have successfully demonstrated that plant pathogenic bacteria can enter a VBNC (viable-but-nonculturable) state following a stress, and resuscitate when the stress is removed and/or in the presence of their host. This is especially the case for the seedborne bacteria *Xanthomonas campestris* pv. *campestris* (Ghezzi et al 1999), *Clavibacter michiganensis* pv. *michiganensis* (Jiang et al 2016) and *P. citrulli* (Kan et al 2019). Different doses of copper sulfate cause the loss of their cultivability, which can be restored when the stress is lifted or in the presence of their host. Furthermore, acidic pH, mimicking the acid treatment of tomato seeds, can induce VBNC state in *C. michiganensis* pv. *michiganensis* (Jiang et al 2016). In all these cases of VBNC state due to intense stresses, the revival of the cell occurs at low levels of bacterial populations sizes. Hence these low populations must be amplified either through the use of highly sensitive detection tools before infections can be revealed or through bacterial growth to be able to cause visible symptoms on host plant (Jiang et al. 2016). In accordance with the existence of a VBNC state, the BLS strains found in the hybrid lots, whose parental lots showed presence of BLS DNA, may have originated from the parental seeds.

Our study reveals VCZ strains in hybrid zucchini seed lots, which may be related to their capacity for floral transmission during pollination. Our seed production monitoring tracked the same parent lots planted in the two different regions. When hybrid seeds from the same cross were infected with VCZ strains, infection consistently occured in the Rhône Valley and never in Limagne. Seed infection *via* the environment could occur through flowers during pollination. Zucchini is an insect-pollinated species whose pollination depends exclusively on bees (Nepi and Pacini 1993). Bees could infect the stigmas and spread VCZ strains within the plots. It has been shown that a honeybee (*Apis mellifera*) carries an average bacterial load of 1 × 10 CFU, which is likely to be deposited on flowers when they are visited (Prado et al. 2020; Ushio et al. 2015; Pattemore et al. 2014). Furthermore, it has been shown that populations of *P. syringae* pv. *syringae* and *P. syringae* pv. *actinidiae* can remain viable in beehives for six to fourteen days, after which bees can spread these bacteria to nearby fields (Pattemore et al. 2014). However, in our survey, the source of VCZ inoculum for the bees was not identified in the samples of weeds, volunteer plants, or zucchini plants that we collected from the plots. In zucchini crops grown for seed production, pollination is carried out by honeybees from hives temporarily placed in the fields. These hives can then be moved to other zucchini fields to ensure pollination. Further analyses of zucchini pollen and bees could clarify the role of bees in the spread of VCZ strains.

The difficulty to identify the sources of inoculum for the VCZ and BLS strains could be due to the small sample sizes that were analyzed for the parental seeds. For example, the seed sample from the male parent (♂ F), whose progeny was contaminated with VCZ and BLS strains, contained only 500 seeds. The other samples analyzed contained approximately 2,000 seeds. This is lower than the 5,000 seeds commonly tested for other diseases such as halo blight on beans caused by *Pseudomonas amygdali* pv. *phaseolicola* (ISTA, 2025) and *P. syringae* pv. *pisi* on pea seeds (ISF, 2020). The sample size analyzed seems small compared to the size of a batch of parental seeds and the number of sown seeds. Indeed, for one hectare of zucchini grown for seed production in open fields, approximately 16,000 male seeds and 40,000 female seeds are sown. In this study, the parental seed lots were distributed across two to four plots ranging in size from 2 to 6 ha. Thus, if we consider that three 4-hectare plots were sown, this represents 192,000 seeds from the male batch and 480,000 seeds from the female batch; therefore, approximately 1.04% of the male batch was analyzed and 0.42% of the female batch (based on an analysis of 2,000 seeds). Given this estimate, it would have been preferable to increase the size of the analyzed samples.

The ability of the BLS strain to transmit through the pericarp seemed to be linked to the ability to cause symptoms on adult plants. This strain could enter through natural openings, such as stomata or lenticels, or through wounds caused during inoculation, while the cuticular layer of *Cucurbita* spp. fruit is still relatively thin (Sutherland and Hallett 1993). In watermelons, it has been shown that *P. citrulli* penetrates the fruit’s pericarp through the stomata up to two or three weeks after anthesis (Frankle et al. 1993). After this period, the stomata on the surface of the watermelon are covered with a waxy layer, and the bacteria can no longer enter through these natural openings (Frankle et al. 1993). The small pustules observed in the area inoculated with the BLS strain could be a sign of the pathogenic interaction between this strain and the plant, which allows it to colonize the fruit and infect the seeds. Bacteria transmitted in this way generally cause symptoms on the fruit before spreading to the seeds, as described for *P. citrulli* and *P. amygdali* pv. *phaseolicola* (Dutta et al. 2015; Taylor et al. 1979). In contrast, the VCZ strain, which does not cause disease in adult plants, did not cause this type of symptom on the pericarp and was not transmitted to the seeds either. As shown for numerous pathogenic bacteria in other organs, the type III secretion system (T3SS) may help these strains manipulate host defenses *via* T3Es, thereby enabling fruit colonization and seed infection.

These results carry a practical implication for controlling seedborne pathogenic bacteria: monitoring should assess transmission through to the seedling stage, not seed infection alone for example, by testing seedlings rather than seeds.

## Supporting information

Suppmentental Figure S2

## Acknowledgments

We thank PHENOTIC Angers for plant production, INRAE (PHENOTIC 2023) and ANAN platform, SFR Quasav for DNA analysis facilities. The authors thank M. Barret for his very fruitful review of the manuscript.

