## Supplemental FIgure S1 for "Insights into the transmission of seedborne *Pseudomonas syringae* strains that cause zucchini diseases"

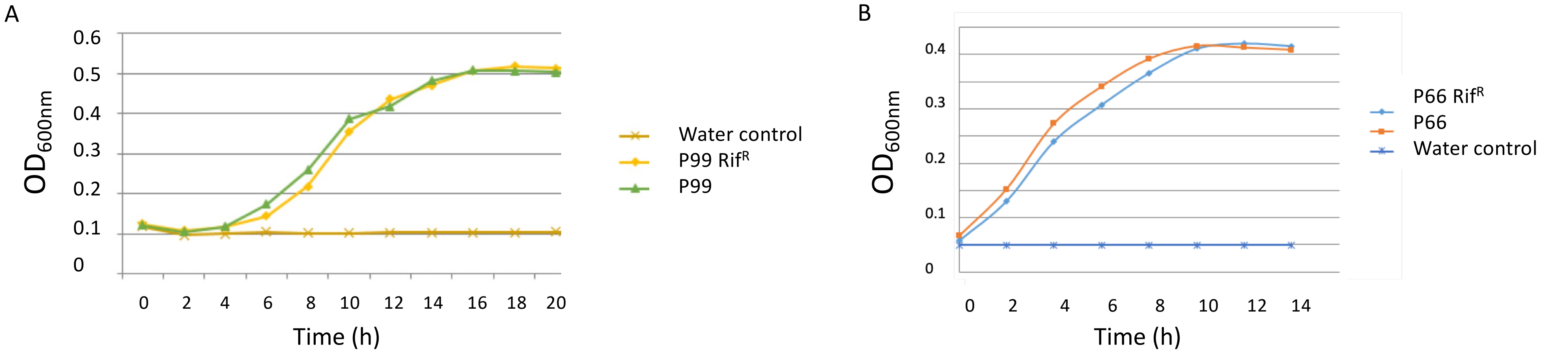


C

| Strains | Stunting | Cotyledon necrosis | vein clearing |
|---|---|---|---|
| **P99** | 8/8 | 8/8 | nd[1] |
| **P99 Rif$^R$** | 8/8 | 8/8 | nd[1] |
| **P66** | 8/8 | 8/8 | 8/8 |
| **P66 Rif$^R$** | 8/8 | 8/8 | 8/8 |
| **Control** | 0/4 | 0/4 | 0/4 |

[1] nd: not determined because there was no first leaf emergence due to complete stunting.

**Supplementary Figure S1. Growth and pathogenicity of the rifamycin resistant variants compared to the wild type strains. A** and **B** Growth curves in 10% TSB of the rifamycin resistant variants P99 Rif$^R$ and P66 Rif$^R$ compared to wild type strains P99 and P66. Three wells per treatment of a 100-well honeycomb microtiter plates (Thermo Electron, France) were filled with 10% TSB (1.7 g/liter tryptone, 0.3 g/liter soybean peptone, 0.25 g/liter glucose, 0.5 g/liter NaCl, and 0.5 g/liter $K_2HPO_4$) and inoculated at a final concentration of $10^7$ CFU/ml. Optical densities at 600 nm were measured every hour using a Bioscreen C instrument (Labsystems, Helsinki, Finland) at 28°C with continuous shaking (120 rpm) until reaching the stationary phase. **C**. Eight plantlets of *Cucurbita pepo* subsp. *pepo* cv. Tosca at the cotyledon stage were inoculated per strain by rubbing the cotyledon surface three times with a finger soaked in bacterial suspensions calibrated at $1 \times 10^8$ CFU/ml. Plantlets were produced in a greenhouse from healthy seeds for 7 days in pots containing Klassman substrate and watered as needed for optimal growth. Inoculated plantlets were incubated in a growth chamber (18°C night, 25°C day, 14 h daylight, relative humidity near 100%), and the health status of each plantlet was recorded seven days after inoculation. Plantlets were qualitatively rated for stunting, cotyledon necrosis and vein clearing on leaves.
