## Supplementary material for "Insights into the transmission of seedborne *Pseudomonas syringae* strains that cause zucchini diseases": Suppmentental Figure S2

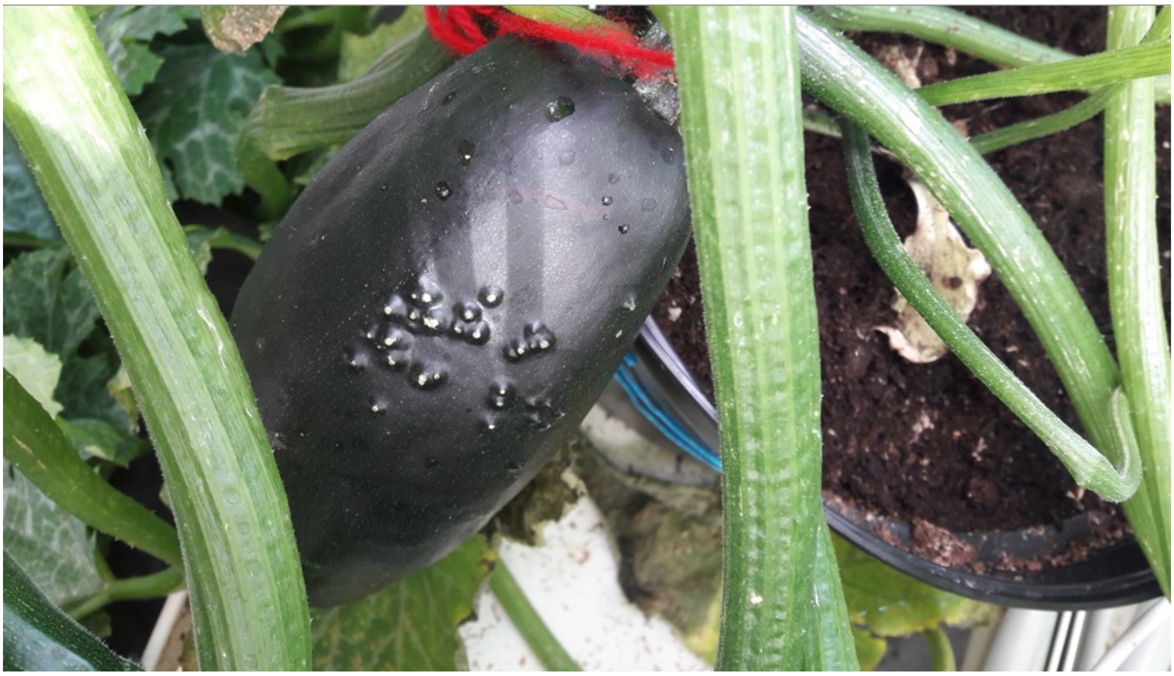

**Supplementary Figure S2. Symptoms observed on fruits inoculated by contact with strain P66 Rif$^{R}$.** One-week-old fruits were inoculated by contact with a 5 × 5 cm square of sterile filter paper soaked with 200 µl of bacterial suspension calibrated at 1 × $10^{7}$ CFU/ml. After removing filter paper, corky pustules were observed in the inoculated area during fruit maturation.
